# Kinetic Machine Learning Unravels Ligand-Directed Conformational Change of μ Opioid Receptor

**DOI:** 10.1101/170886

**Authors:** Evan N. Feinberg, Amir B. Farimani, Carlos X. Hernandez, Vijay S. Pande

**Author notes:** Denotes equal contribution (Evan N. Feinberg, Amir B. Farimani).

## Abstract

The μ Opioid Receptor (μOR) is a G-Protein Coupled Receptor (GPCR) that mediates pain and is a key target for clinically administered analgesics. The current generation of prescribed opiates – drugs that bind to μOR – engender dangerous side effects such as respiratory depression and addiction in part by stabilizing off-target conformations of the receptor. To determine both the key conformations of μOR to atomic resolution as well as the transitions between them, long timescale molecular dynamics (MD) simulations were conducted and analyzed. These simulations predict new and potentially druggable metastable states that have not been observed by crystallography. We applied cutting edge algorithms (e.g., tICA and Transfer Entropy) to guide our analysis and distill the key events and conformations from simulation, presenting a transferrable and systematic analysis scheme. Our approach provides a complete, predictive model of the dynamics, structure of states, and structure–ligand relationships of μOR with broad applicability to GPCR biophysics and medicinal chemistry.

## Introduction

Over the past decade, an epidemic of opioid abuse has increasingly plagued the United States^1^. This health crisis is catalyzed in part by the elevated administration of opioid therapeutics to treat acute and chronic pain. Since the solution of the first structure, crystallography of GPCRs has both illuminated the structural biology and empowered medicinal chemistry of this class of receptors^2,3^. Recently, crystal structures of μOR itself were solved in its “inactive” and “active” conformations^4,5^. However, other biophysical and pharmacological experiments have definitively demonstrated that μOR traverses multiple functionally important conformational states^6–8^.

In this paper, we endeavor to (1) discover states of μOR that have thus far been refractory to crystallography and (2) juxtapose how opiates of different scaffold classes tune the receptor toward distinct conformational energy landscapes. To achieve these aims, we conducted unbiased, atomistic molecular dynamics (MD) simulation of μOR for 250 μs in each of three conditions: unliganded, BU72^9^-bound, and Sufentanil^10^-bound. The former ligand is a higher affinity derivative of Buprenorphine, a drug administered to treat opioid addiction and acute pain, while the latter is a higher affinity derivative of Fentanyl, a drug administered as a component of operative anesthesia.

Both building on and departing from preceding GPCR simulation works^11,12^, we undertake a systematic statistical approach to analyze this vast simulation dataset. In particular, we discover the slowest conformational degrees of freedom accessible to μOR with the time-independent component analysis (tICA) algorithm ^13–15^, construct Markov State Models^16,17^ (MSM) based on these tICA degrees of freedom to compare μOR’s conformational dynamics between conditions, and finally postulate causal linkages between events using transfer entropy^18,19^ analysis. The tICA and Transfer Entropy approaches define two related measures of importance for each amino acid residue in μOR. By measuring slowness, tICA identifies groups of amino acid residues whose transitions constitute the highest kinetic barriers to conformational change in the system of interest. On the other hand, computing the PageRank^20^ of each inter-residue contact in the mutual transfer entropy network measures the causal centrality of each such contact in triggering conformational change in the receptor. We apply both tICA and transfer entropy in this manner to determine two complementary measures of “importance” for each inter-residue contact (cf. Methods for further details).

We note that this systematic approach should be transferrable to other conformationally plastic proteins which experience allosteric modulation, such as other GPCRs, ion channels, and nuclear receptors. Specific to μOR, stark thermodynamic differences are observed between the behavior of BU72 and of Sufentanil. The statistical models reveal allosteric modulation pathways that connect perturbations in the ligand binding pocket with global conformational changes of the receptor.

## Results

### I. Rearrangement of D147^3.32^, F289^6.44^, W293^6.48^, and Y326^7.43^ gates activation

The experimental and computational literature of GPCR conformational dynamics, which include paired inactive and active state crystal structures (e.g., refs ^4,21–23^); NMR, EPR, and DEER studies (e.g., refs.^8,24,25^), mutagenesis, and MD simulation (e.g., refs ^11,12^), have indicated common features of GPCR activation, including an outward movement of transmembrane helix 6 (TM6) upon activation^23,26^, the importance of a salt bridge (the “ionic lock”) between charged residues on TM3 and TM6, and the rearrangement of the conserved NPxxY domain (residues 7.48-7.53)^26^. In contrast to B2AR, Rhodopsin, and many other GPCRs, μOR features a hydrogen bond between R165^3.50^ and T279^6.34^ which plays a role in mediating activation according to mutagenesis and molecular dynamics studies.^21,27^ Previous long-timescale MD simulations of B2AR^11,12^ and of μOR^5^ have suggested that the conserved residues I^3.40^, P^5.50^, F^6.44^ and W^6.48^ play key roles in allosteric coupling between the binding pocket and intracellular conformational changes.

By analyzing both the kinetic slowness of inter-residue contacts with tICA as well as their PageRank in the transfer entropy network of μOR (cf. methods for more details), we determined the critical signaling motifs of the receptor. Both metrics identify a quartet of residues – D147^3.32^, F289^6.44^, W293^6.48^, and Y326^7.43^, which we term **DFWY** – as a critical gate of activation (Figure 1, E.D. Table 5, E.D. Figure 9). The significance of this DFWY motif is reflected both in its slow kinetics as well as its centrality within the causality graph (as described in Methods section). In addition, receptor deactivation, in which μOR transitions from its active crystal-like structure toward either an inactive crystal-like or other terminally inactive state, is triggered by reconfigurations of these residues. First, D147^3.32^ and Y326^7.43^ in the binding pocket move apart by 3 Å from each other and therefore also from the center of the binding pocket).

**Figure 1:**
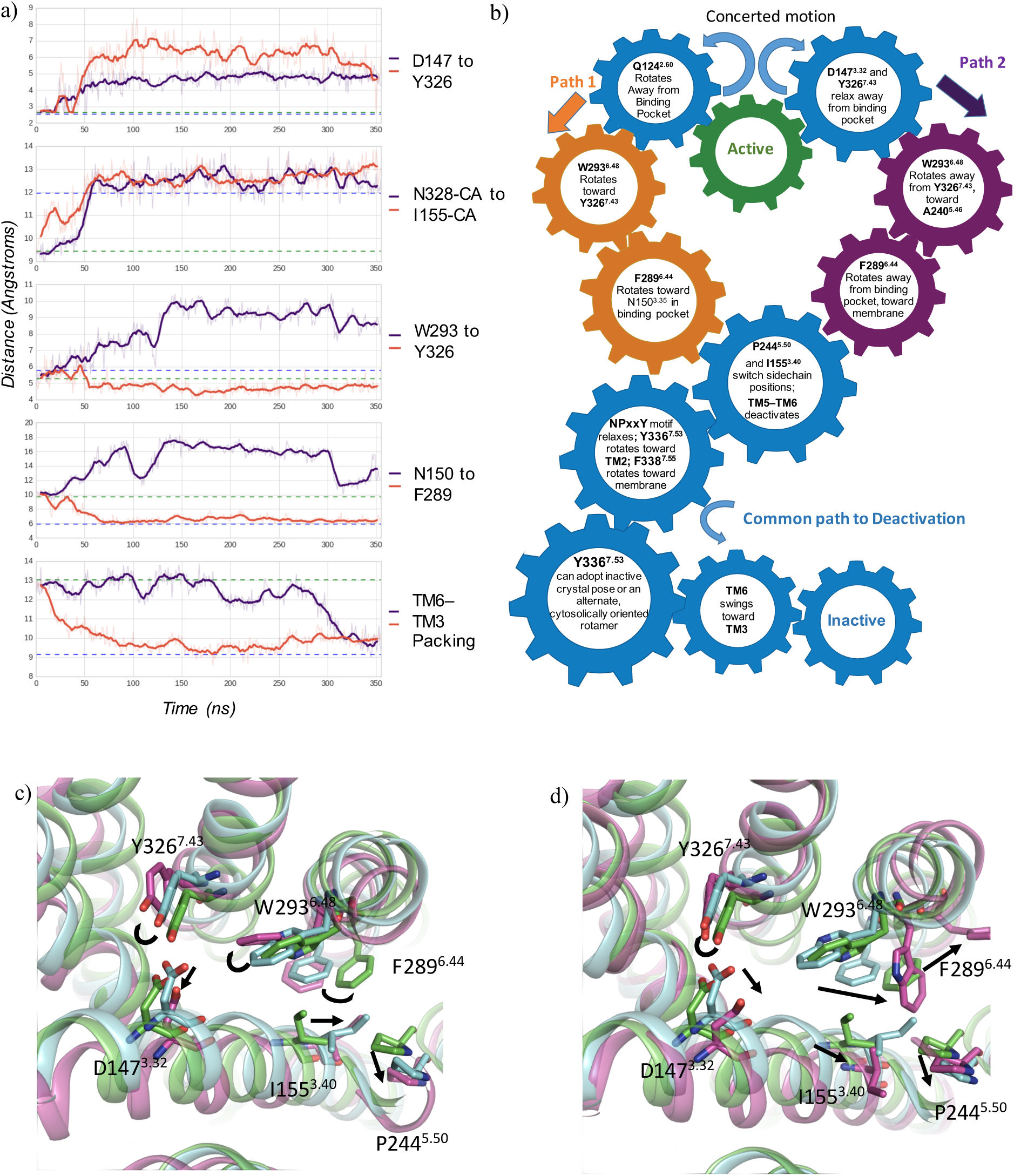
μOR samples at least two deactivating pathways that are gated by two distinct configurations of the DFWY motif, (a), (b) and (c): “Canonical” pathway (orange trace) entails (1) D147^3.32^ relaxation away from Y326^7.43^, (2) W293^6.48^ rotation toward Y326^7.43^, (3) F289^6.44^ rotation toward binding pocket to maintain 7r-stacking with W293^6.48^, (4) I155^3.40^ switches to take space previously occupied by F289^6.44^, (5) P244^5.50^ relaxes away from 1155^3.40^ toward inactive position, triggering deactivation of TM6-TM5 packing and ultimately the relaxation of TM6 toward its inactive pose occluding the G Protein coupling domain, (b) and (d): In the alternative pathway, W293^6.48^ instead rotates toward TM5 and F289^6.44^ rotates outward toward the membrane. In (c) and (d), green **→**1 active crystal structure (PDB: 5C1M), cyan inactive crystal (PDB: 4DKL), and purple a snapshot from MD simulation (different frames for (c) and (d)).

This loss of hydrogen bond between the negative carboxylic acid of D147^3.32^ and the phenolic hydroxyl of Y326^7.43^ represents a larger transition between “pinched” active-like and “relaxed” inactive-like binding pocket configurations (Figure 1).

The computed tICA and transfer entropy models underscore the significance of transitions of Y326^7.43^ and W293^6.48^. These two residues trigger signaling cascades that propagate from the binding pocket toward the cytosol. Once the D147–Y326 hydrogen bond is broken, the sidechain of W293^6.48^ can trigger two distinct modes of deactivation. In one pathway ‐‐ which we term “canonical” due to its parallels with B2AR ‐‐ W293^6.48^ rotates away from A240^5.46^ and F241^5.47^ on transmembrane helix 5 (TM5) and toward both G325^7.42^ and Y326^7.43^ on TM6. Conversely, W293^6.48^ can instead rotate away from Y326^7.43^ and toward the carbonyl oxygen of A240^5.46^ (Fig. 1 and Fig. 4).

In turn, these two possible W293^6.48^ transitions enable two distinct modes of relaxation of F289^6.44^ away from P244^5.50^ (Fig. 1). Along the “canonical” pathway, F289^6.44^ rotates toward TM3 and the receptor core (as measured by the N150^3.35^–F289^6.44^ distance, cf. Fig. 1 and E.D. Fig. 1). In contrast, in the alternate pathway, F289^6.44^ rotates away from the receptor and toward the membrane (Fig. 1b and 1d; E.D. Fig. 1). The coupling between F289^6.44^ and W293^6.48^ likely stems in large part from the stable π-π stacking between the aromatic sidechains of the two residues. The loss of a hydrophobic contact between F289^6.44^ and P244^5.50^ in turn allows TM5 to transition from its bulged, active-like configuration and toward its relaxed, inactive-like configuration.^i^

Ultimately, the signal stemming from the DFWY rearrangement is transduced to the signaling termini – the G Protein coupling domains ‐‐ of the receptor. Triggered by the rotation of Y326^7.43^ toward TM2 and, in the canonical pathway, stabilized by the combined rotation of W293^6.48^ and F289^6.44^ toward Y326^7.43^, N328^7.45^ translates up to 3 Angstroms away from TM6 and from I155^3.40^ on TM3. The shift of N328^7.45^ is correlated (Fig. 1) with the highly sequence conserved Y336^7.53^ in the NPxxY (residues 7.49–7.53) motif^11,28^, which undergoes a critical transition from its bulged, TM6–oriented position toward its relaxed, TM2–oriented rotamer. Therefore, this overlapping quartet of– I155^3.40^, P244^5.50^, F289^6.44^, and N328^7.45^ – is a key mediator of activation whose rearrangement influences both the overall packing of TM5 against TM6 as well as the NPxxY relaxed vs. bulged configuration.

Intriguingly, while *apo* (i.e., unliganded) and BU72-bound simulations traverse both pathways to deactivation, Sufentanil-bound simulations overall less frequently and are biased toward the alternate, W293–Y326 far / W293–A240 close pathway (E.D. Fig. 1, rows 3 and 4). Specifically, the unliganded μOR samples the inactive state in 2.9% of all frames (cf. Methods for definition of inactive), BU72-bound μOR in 3.7% of all frames, and Sufentanil-bound μOR in 0.6% of all frames, a several fold difference. Among inactive conformations sampled by MD, the unliganded μOR features a W293–Y326 far / W293–A240 close configuration in 9.0% of all frames, the BU72-bound μOR in 6.0% of all frames, and the Sufentanil-bound μOR in 65.2% of all frames, an order of magnitude difference.

While previous studies^11,12,21,27^ have underscored the TM3-TM6 distance in general and the polar contact between the DRY motif on TM3 and residues on TM6 as a critical gate for the rearrangement of these helices, the simulation dataset presented here suggests the R165^3.50^– T279^6.34^ hydrogen bond as only one of several key components of μOR dynamics. The Sparse tICA algorithm (E.D. Figure 10) identifies R165-T279 as a component of the eighth-slowest degree of freedom of the receptor. Meanwhile, the rearrangement of D147, W293, and Y326 near the binding pocket as well as F289, P244, N328, and I155^3.40^ in the interceding region between the binding pocket and cytosol (discussed in ref^5^) are components of the second-, fourth- and fifth slowest degrees of freedom, indicating that the rearrangements discussed in this section are a necessary, slower step that enables the ultimate TM3-TM6 activation necessary for G Protein coupling.

### II. Ligands induce novel transitions in both the binding pocket and G Protein coupling domains that define functionally diverse conformational states

The residues described in Section (I) can be depicted as key molecular switches, the toggling of which defines a series of intermediate macrostates that the receptor traverses between its inactive and active crystal-like conformations. However, these residues (as well as others to be described) also define global receptor conformations that are orthogonal to reaction coordinates that connect the two crystal-like states. These new conformers are available to the reader as PDB files in the supplementary information section.

The tICA algorithm identifies the slowest collective degrees of freedom of a biomolecular system, and these slow motions correspond to the biologically important dynamics of the system. The slowest tICA reaction coordinate, which we denote ω_1_, involves rearrangements of the binding pocket and extracellular regions of TM5, TM6, and TM7 that define receptor states distinct from either crystal structure. Specifically, K233^5.39^, K303^6.58^, and W318^7.35^ adopt multiple configurations. The importance of these residues is concordant with several experimental studies, which have identified, for example, the role of K233^5.39^ and W318^7.35^ in conferring ligand agonism and of K303^6.58^ in conformational change^4,24,29,30^.

While some ω_1_ states are accessible to μOR regardless of simulation condition (BU72, Sufentanil, or *apo*), it is clear that the Sufentanil induces the μOR to sample ligand-dependent metastable states (Fig. 2). One such state entails (a) a 4.0 Å shift of W318^7.35^, and much of the extracellular half of TM7, toward TM2; (b) a partial unfolding of the extracellular helix of TM6, enabling K303^6.58^ to rotate toward E229; (c) a rotamer shift of Y336^7.53^ toward F289^6.44^, forming a contact between the phenolic hydroxyl of the former and backbone amide of the latter. Since TM6 adopts its outward position and Y336^7.53^ shifts up to 1.0 Å closer to F289^6.48^ than it does in the active crystal, we define this Sufentanil-induced state as active-like, and denote it R^∗-SUF^.

**Figure 2:**
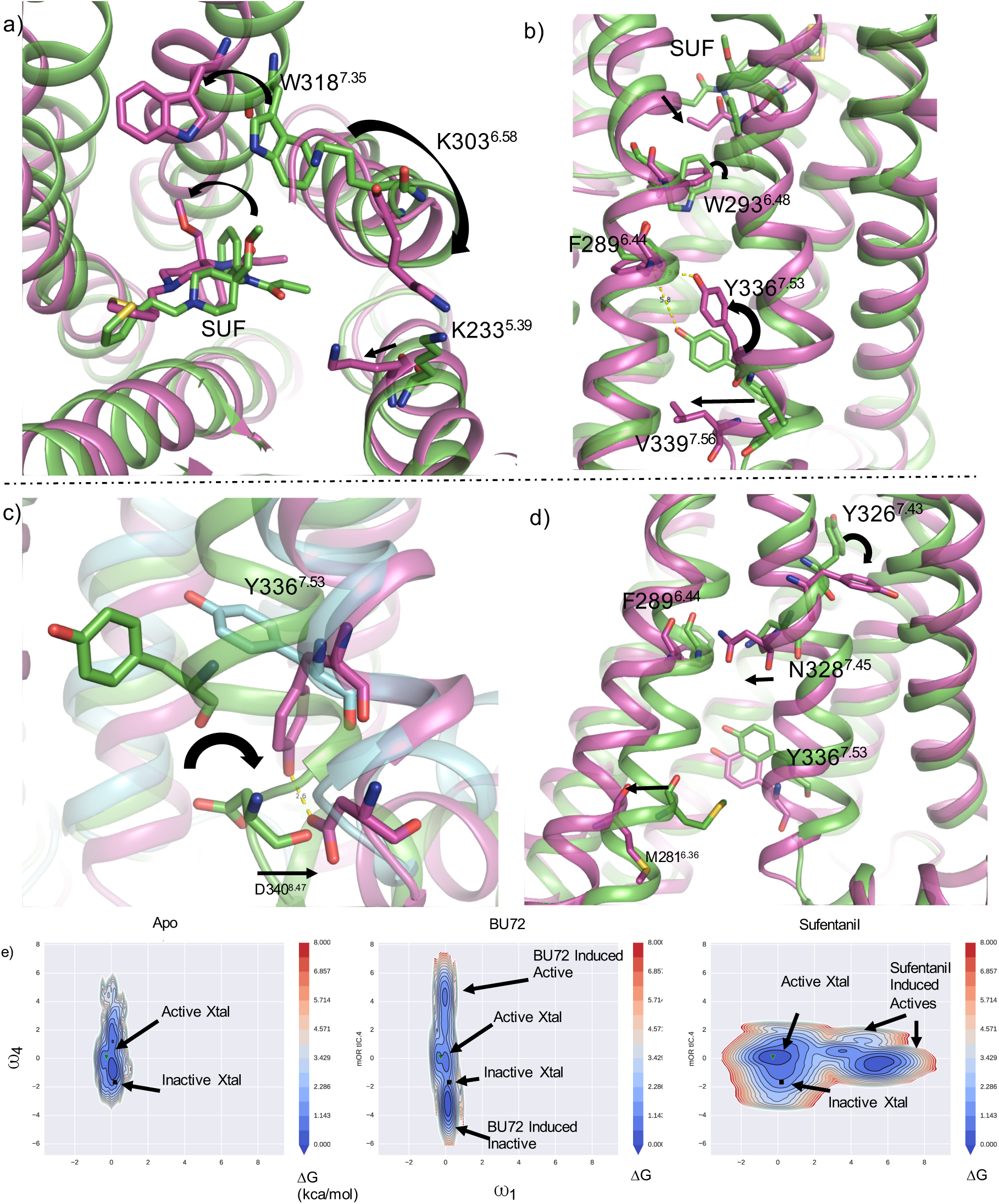
Non-canonical states of μOR. (a)-(d): Active crystal in green, inactive crystal in cyan, MD snapshot in purple, (a) and (b) Sufentanil-induced,high ω_1_, active-like macrostate of μOR. (c) BU72-induced, low ω_4_, inactive-like macrostate of μOR. d) BU72-induced, high ω_4_, active-like macrostate of μOR. e) Markov State Model reweighted free energy plots (kcal/mol) of μOR projected onto tICA coordinates ω_1_ and ω_4_ in three different conditions: from left to right, Apo, BU72, and Sufentanil.

In summary, ω_1_ defines a reaction coordinate that distinguishes Sufentanil-bound μOR from BU72-bound and unliganded μOR in conformational space. Analogously, ω_4_ and ω_5_, the fourth and fifth slowest tICA coordinates, define BU72-induced states that are distinct from either crystal structure and sampled substantially less frequently by Sufentanil-bound or unliganded μOR (cf. E.D. Table 11 for relative tICA timescales). One such macrostate entails a novel rotamer of Y336^7.53^ oriented toward the cytosol, where it is stabilized by hydrogen bonding with D340^H8^ (Fig. 2c).

Furthermore, our modeling predicts that two neighboring salt bridges – R277^6.32^-D340^H8^ and R280^6.35^-E341^H8^ – may play key roles in gating activation (Fig. 3, Movie 1, E.D. Figure 14). Particularly, formation of the R280-E341 interaction breaks the salt bridge that E341^H8^ forms with K344^H8^ in both crystal structures. Notably, a transition centered on K344^H8^ is implicated as a key initial component in receptor activation in recent experimental studies^24^. Complementarily, mutations in E341^H8^ have been shown to significantly modulate the constitutive activity of μOR^31,32^. It is therefore reasonable to hypothesize that this ω_4_ high state (Fig. 2c, Fig. 3) precedes the inactive crystal structure on the overall activation pathway. Specifically, we propose that the R280-E341 salt bridge complements the more sequence conserved DRY motif^33^ R165-T279 hydrogen bond in polar mechanisms of gating activation.

**Figure 3:**
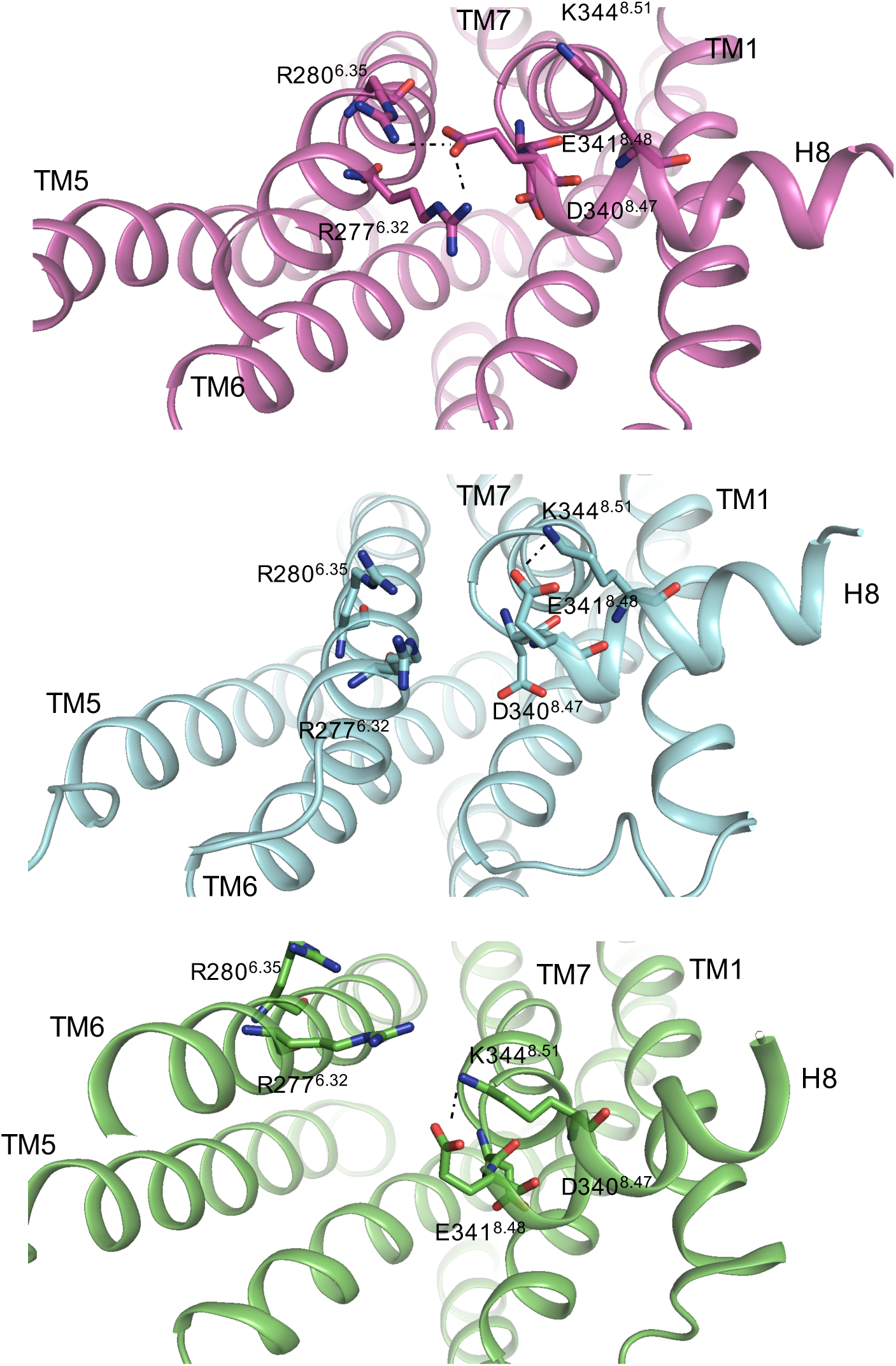
Inactive-like structure captured from MD simulation shows non-crystallographic salt bridge forming concomitantly between both R280^6.35^ and R277^6.32^ with E341^8.48^. Purple: MD snapshot; Cyan: Inactive crystal (PDB: 2RH1); Green: Active Crystal (PDB: 3P0G). This MD snapshot is the same as that displayed in main text Figure 3c, with a different orientation and different residues highlighted. In both inactive and active crystal structures, E34 1^8.48^ instead forms a salt bridge with K344^8.51^ yet interacts neither with R280^8.35^ nor R277^8.32^. Note, as displayed in main text Fig 3c, that this structure also involves a polar interaction between D340^8.47^and Y336^7.53^. Due to the stability of this state (cf. E.D. Figure 1, Row 6), and the proximity of this rearrangement to the G Protein binding domain, this state maybe biologically important. In addition, it is notable thatrecent experimental work has revealed the significance of early changes in H8 (measured by K344^8.51^ probe in particular) in propagating initial steps of μOR conformational change^1^. Therefore, we propose thatthis state represents the *initial* inactive structure along the activation pathway.

In the BU72-induced, ω_4_-high state, Y326^7.43^ adopts a third position oriented toward C79^1.43^, effectively pushing neighboring helical turns of TM7 by at least 2.0 Å toward TM6. In turn, an interaction of N328^7.45^ induces a bend near hinge residue F289^6.44^, stabilizing TM6 in an alternate active-like conformation (Fig. 2d). The proximity of Y336^7.53^ and of the intracellular portion of TM6 to the putative G Protein and Arrestin binding domain suggest the potential biological significance of these states.

### III. BU72 transitions either to a beta–FNA-like deactivating pose or to a series of non-crystallographic positions buried in the receptor core

In nearly 250 μs of MD simulation, BU72 samples several metastable poses that are in turn coupled to conformational states of the receptor as a whole. Transfer entropy analysis of a network of μOR-BU72 contacts and μOR tICA components (cf. Methods for details) indicates that binding pocket residues W293^6.48^, H297^6.52^, Q124^2.60^, W133^ECL1^, Y326^7.43^, and M151^3.36^ are particularly important in modulating the receptor’s state (E.D. Figure 6 and E.D. Tables 1, 3, and 4).

In the canonical deactivation pathway, the morphinan scaffold of BU72 transitions from its active, crystal-like pose toward a pose that is analogous to beta-Funaltrexamine’s inactive crystal pose (Fig. 4c, E.D. Fig. 2, E.D. Movie 1). The high PageRank of Q124^2.60^ in the transfer entropy network of μOR underscore its centrality in a polar network that rearranges upon receptor deactivation (Figure 4; E.D. Figure 6, Row 2, Column 1; E.D. Tables 1 & 2). Note how, in the active crystal, the non-morphinan phenyl ring of BU72 contacts W133^ECL1^ and Q124^2.60^, while Q124^2.60^ hydrogen bonds with Y326^7.43^ (Fig. 4c). The rotation and translation of BU72 toward TM5 as it transitions from its canonical active to inactive pose initiates a cascade of events in simulation: (1) Y326^7.43^ retreats toward TM2; (2) Q124^2.60^ rotates extracellularly to engage with the W133^ECL1^ indole; (3) TM2 as a whole, including Y128^2.64^, relaxes away from the binding pocket and toward the membrane; and (4) H319^7.36^ loses its interaction with Y128^2.64^, freeing it to rotate with its aromatic partner W318^7.35^ and the rest of TM7 as a nearly rigid body toward TM6.

**Figure 4:**
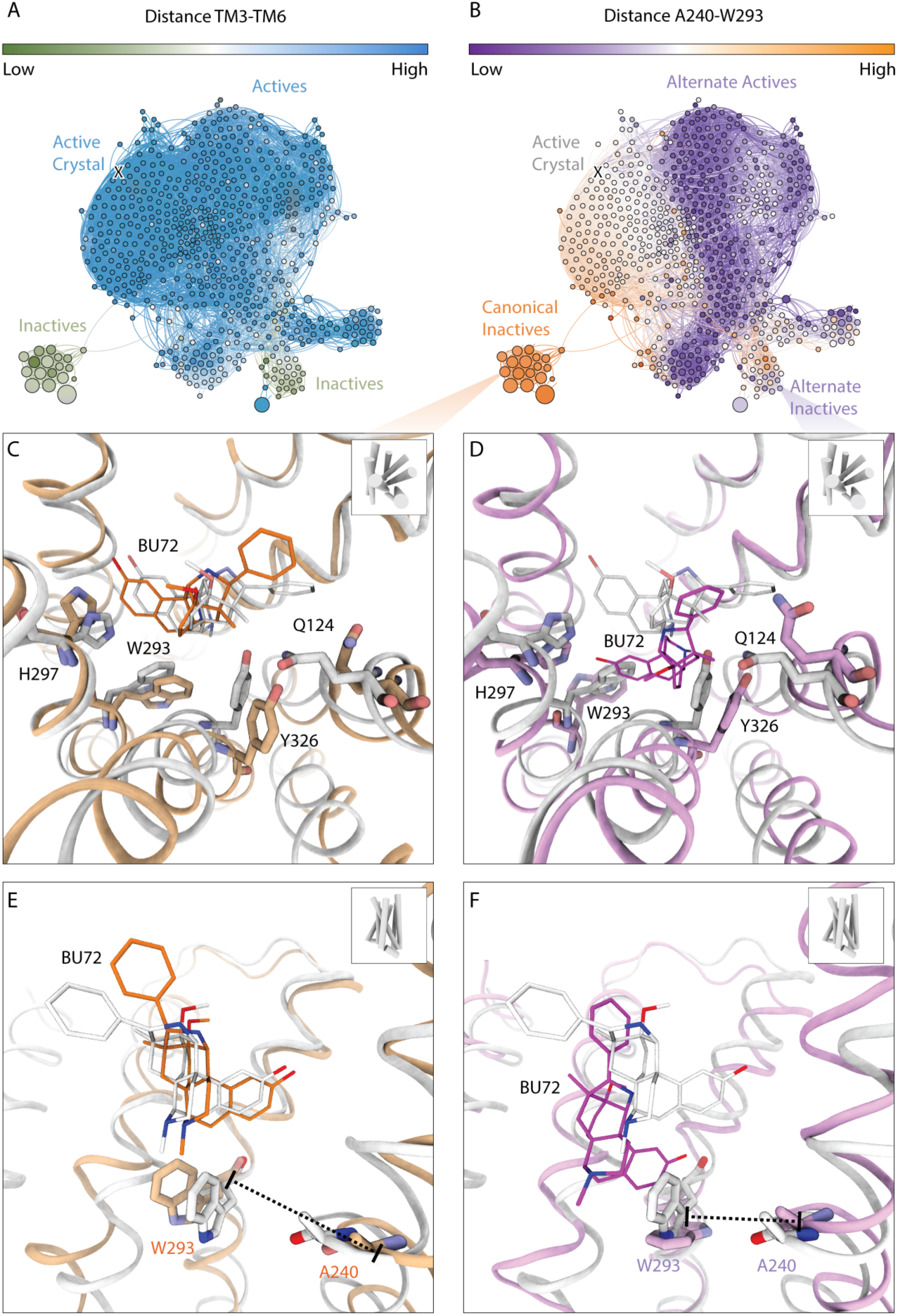
BU72 samples distinct deactivating poses which trigger distinct deactivation pathways of μOR. (a) and (b): Network views of Markov State Model (MSM) of BU72-bound μOR. Each node is an MSM state, each edge is a possible transition, Nodes are colored by TM6–TM3 distance in (a) and by A240-W293 distance in (b). Nodes are sized proportional to MSM equilibrium populations, In c-f, active crystal displayed in grey, MD snapshot displayed in orange for canonical pathway (c, e) and purple for alternate pathway (d, f). C and E) Canonical deactivation in presence of BU72 in binding pocket. Particularly noteworthy changes in the binding pocket include the rotamer shift of Q124 away from Y326, the rotation fo Y326 toward TM2, the rotation of W293 toward Y326, and H297’s rotation toward TM5. D and F) Alternate deactivation where BU72 plunges several angstroms into receptor core, inducing non-crystallographic rotamers of W293 and, therefore, F289. Notably, the rotation of Q124 away from Y326 and of Y326 toward TM2 is common to both pathways.

In addition to transitions between the crystallographic opiate poses, in MD simulation BU72 also populates several orientations that either induce or stabilize μOR to occupy similarly noncrystallographic conformational states. Several such poses sample regions of phase space that would be forbidden by most, if not all, docking techniques. The slowest transition of BU72 as measured by tICA entails an at least 6.0 Å and up to 12.0 Å translation and rotation in the cytosolic direction, probing into the core of the receptor (Fig. 4d, 4f, Movie 2). This cytosolic transition is defined by a rotation of the BU72 ether toward the hydrophobic region comprised of Ile296, Cys321, Ile322; a novel polar interaction forming between the BU72 phenol hydroxyl, a functional group conserved across many opiates, and H297^6.52^; and 140° χ^2^ rotamer shift of W293^6.48^ (akin to that observed in ref^5^), leading to a fully connected aromatic network involving W293^6.48^, H297^6.52^, and the BU72 phenol. In another pose, in which the morphinan phenol rotates 6.0 Å to pack against A240^5.46^ and the non-morphinan phenyl stabilizes Q124^2.60^ in an inward position, BU72 stabilizes the high ω_4_ active state described in the previous section (E.D. Figure 1 row 5 column 2; E.D. Figure 10b). A third distinct pose entails (a) a 2.0 Å closer contact between the BU72 phenyl ring and the indole of W133^ECL1^, and (b) a closer packing of the aliphatic bridge region of BU72 with Y326^7.43^ (E.D. Figure 1a, row 3, column 1; E.D. Figure 7a). Since all three poses are distinct from the crystal structures and sampled in the presence of and/or stabilize cytosolically active conformations of μOR, it raises the possibility that the distinctly high affinity and agonism of BU72 is in part conferred by its configurational entropy.

### IV. Sufentanil is highly stable in its modeled active pose and drives state transitions by exchanging its contacts with Q124^2.60^, W133^ECL1^, W293^6.48^, and Y326^7.43^

Fentanyl and its analogs are high affinity agonists often used in anesthesia and pain control, but existing μOR crystal structures contain only morphinan derivatives. In lieu of crystallography, we predicted the binding pose of Sufentanil by using docking to obtain potential starting points and then deploying MD to identify and optimize the correct pose (Fig. 5; Methods; S.I. PDB file). Intriguingly, as aforementioned, the Sufentanil-bound μOR deactivated less frequently than in the unliganded or in the BU72 bound simulations (E.D. Figure 1).

**Figure 5:**
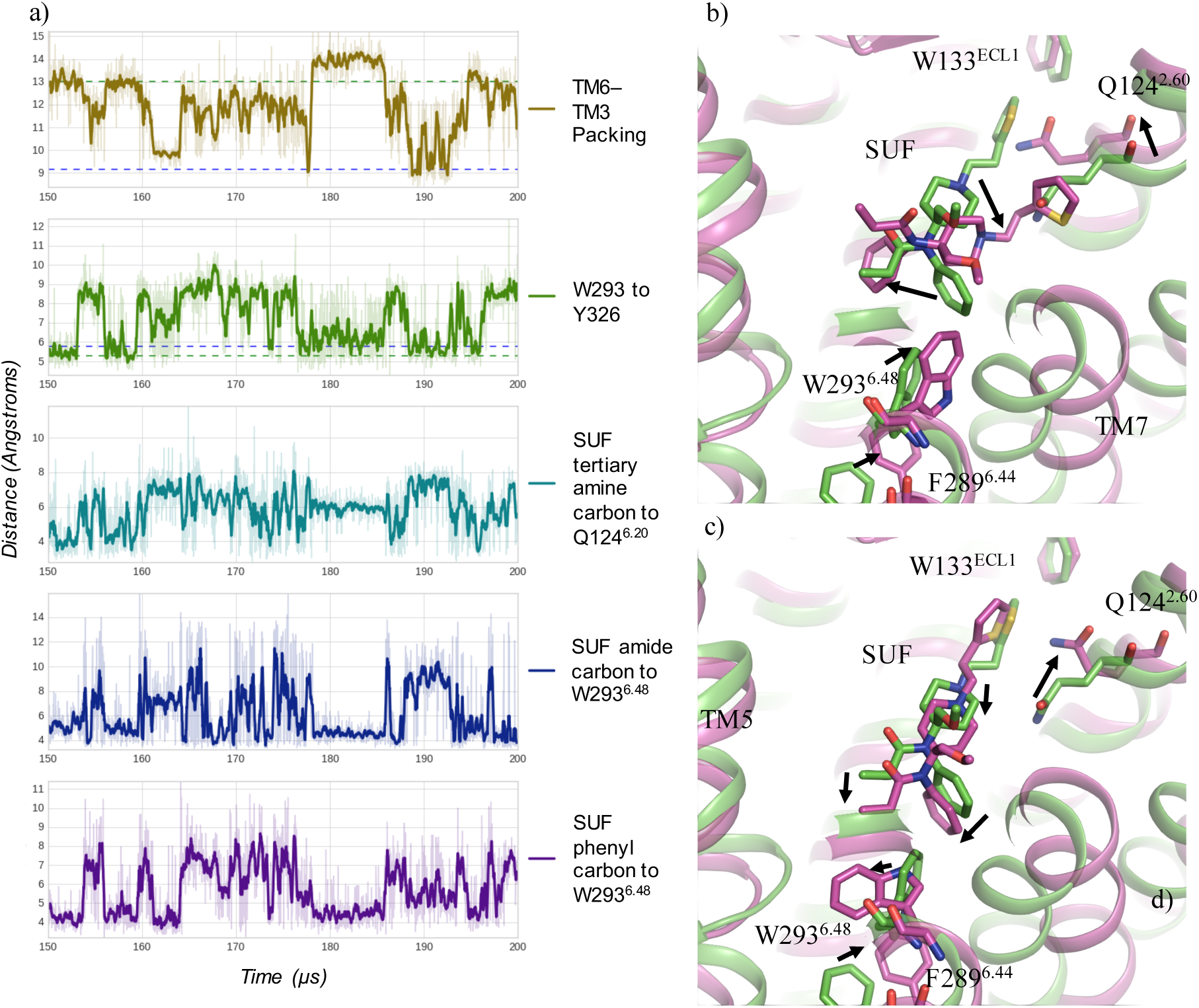
Sufentanil samples several deactivating poses, all of which are predicated on transitions of its aromatic rings with respect to Q124^2.60^ and W293^6.48^ a) Markov State Model representative timeseries in which μOR undergoes distinct deactivation events. (a) and (b) At ~190 μs mark, a several angstrom movement of Sufentanil phenyl with respect to W293^6.48^ is a key factor in the canonical pathway of relaxation of TM6. Concurrently, Sufentanil methoxy oxygen shifts to contact the W293^6.48^ indole, and Q124^2.60^ exchanges its position with the Sufentanil thienyl ring. (a) and (c) At ~160 μs mark, loss of contact between the protonated tertiary amine nitrogen of SUF and the amide sidechain of Q124^2.60^ leads to deactivation as well. Along this non-canonical deactivation path, Sufentanil shifts 1-2 Angstroms into the receptor core, inducing W293 to rotate toward A240. This is also mediated by a Q124 rotamer shift toward the extracellular region. In (b) and (c), green corresponds to docked and MD-optimized pose of Sufentanil in the active crystal structure, while purple corresponds to a different MD snapshot for (b) and for (c).

Simulation suggests that the protonated tertiary amine nitrogen comprises a critical structural motif that not only confers Sufentanil’s affinity for μOR but also contributes to its agonistic character as well. Specifically, the positively charged nitrogen attracts not only D147 – a common motif among the vast majority of known small molecule opiates – but also the carbonyl oxygen of Q124^2.60^, further stabilizing Y326^7.43^’s active position via hydrogen bonding (E.D. Figure 11). In contrast to morphinan and buprenorphine derivatives, like beta-Funaltrexamine and BU72, Sufentanil interacts with Q124^2.60^ directly with the protonated amine present in nearly all non-endogenous opiates, since this latter motif is predicted to rest in the binding pocket 3.7 Å closer to the extracellular region than it does in BU72.

The rotamer of key switch residue W293^6.48^ is controlled by both Sufentanil’s amide and phenyl substituents. In the predicted active pose, the Sufentanil phenyl π-stacks both with Y326^7.43^ and W293^6.48^, stabilizing these two DFWY residues in their active configuration. Conversely, when Q124^2.60^ rotates extracellularly and exchanges its contact with the Sufentanil tertiary nitrogen for the Sufentanil thienyl ring, the ligand adopts one of two poses that can trigger deactivation (Fig. 5). In addition, Sufentanil can undergo a rotation in which the Sufentanil phenyl contacts TM2 and TM3, in turn stabilizing the ω_1_–high state discussed in the previous section (Fig. 2a, E.D. Figure 4c).

## Discussion

This paper outlines the key conformational changes accessible to μOR in the absence of ligand and in the presence of morphinan and fentanyl derivatives. The combined application of tICA and Markov State Models to our MD simulation dataset indicate that the bound ligands – BU72 and Sufentanil – have a marked impact on the conformational energy landscape of the receptor. These two high affinity agonists modulate the configuration of a common set of residues in distinct ways to achieve their unique downstream effects on μOR. Furthermore, μOR deactivated relatively infrequently in the presence of Sufentanil compared to both unliganded and BU72-bound conditions; the added stability conferred to active states of the receptor corroborates the predicted pose of Sufentanil.

This statistical framework also leads to several experimentally testable hypotheses for biophysicists and molecular biologists. For example, mutation of binding pocket residues Q124^2.60^, W133^ECL1^, H297^6.6.52^, Y326^7.43^, W293^6.48^, should all significantly impact the agonism of both BU72 and of Sufentanil. Conversely, mutating D340^H8^ could destabilize one of two deactivating rotamers of Y336^7.53^, thereby stabilizing the active state of the receptor. Similarly, introducing a disulfide bridge between the extracellular domains of TM1 and TM7 could prevent the rigid body rotation of TM7 toward TM6 as well as the rotation of TM1 to the membrane, thereby potentially serving as an activating mutation.

With the great power afforded by MD simulation inevitably comes the high dimensionality and sheer number of data points that renders it so difficult to analyze in a practicable manner. By judiciously applying the tICA algorithm, Markov State Models, and transfer entropy, it is our goal to have produced a replicable framework that might enable other labs to gain actionable knowledge about their protein systems of interest through the “computational microscope”^34^ that is MD simulation. While this paper focuses on biophysics insights derived from unsupervised machine learning analysis of large-scale MD simulations, we have recently shown that the alternative conformations derived from these techniques can also motivate a successful rational drug discovery campaign for μOR that identified a fundamentally novel active chemotype for the receptor.^35^ To facilitate further biophysical and pharmacological analysis, we publish our simulations – both raw and featurized trajectories – as an open source, downloadable resource, useful to both the opioid field in particular as well as the wider structural biology and medicinal chemistry communities.

## Methods

The GPCR-specific Ballesteros-Weinstein scheme^36^ was utilized to denote amino acid residues in the superscript.

To setup simulations, the RCSB PDB^37^ versions of PDB IDs 5C1M and 4DKL – the crystal structures of μOR – were downloaded. Subsequently, after deleting atoms from residues Gly52-Ser64 from 5C1M, and deleting the nanobody as well, the missing residues Arg348-Ile352 were modeled into 5C1M based on the positions of the same residues in 4DKL. To complete side chains and ascertain protonation states at physiological pH, Schrödinger Maestro Protein Prep Wizard was applied. A custom TCL code was then applied to embed the active crystal structure in a POPC bilayer, align it to the orientation described in Orientations of Proteins in Membranes Database, solvate the entire system with TIP3 waters and 0.15 mM of NaCl, and generate a CHARMM36 compatible PSF file by calling the PSFGen program. The Chamber function in ParmEd was then used to convert the resultant PDB and PSF file into AMBER compatible prmtop and inpcrd files, and to assign CHARMM36 force field parameters^38,39^. Finally, simulations were initiated with 2.5 fs in Amber14 on the XStream GPU cluster using the CUDA-enabled variant of PMEMD. The sequence of heating and equilibration was modeled on that described in ref^21^. A total of 1,778 trajectories were generated with combined simulation time of 755.979 μs, summarized in Extended Data Table 11.

The Conformation software package^40^ was written for and applied to the featurization of this large GPCR MD dataset. To summarize, all residue-residue pairs within 6.6 Angstroms measured by closest heavy atom distance in either crystal structure were selected. Then, for each of these approximately 2,200 residue-residue pairs, both the closest heavy atom distance and Calpha distance was computed for each frame in each trajectory, leading to 4,400 “features” for each trajectory frame. Next, the Sparse tICA algorithm was applied to determine the reaction coordinates, or slowest collective degrees of freedom (up to the ten slowest in this case), of the protein. Penultimately, a K-Means model was trained with K=1,000 clusters. Of vital importance for the reader and practitioner to note, this featurization, tICA, and K-Means model was applied equally to all three simulation conditions. Therefore, the unliganded, BU72-bound, and Sufentanil-bound conditions could be equitably compared since they are examined in the same continuous (tICA) and discrete (K-Means) spaces. Finally, a Markov State Model (MSM) was constructed with lag time 25 ns and prior counts 1×10^−5^. The equilibrium state probabilities from the MSMs were used individually in each condition to generate the free energy surfaces projected onto the features and tICA coordinates displayed in this manuscript.

To assess the ability of ligand configuration to influence the conformational state of the receptor, for simulations with Sufentanil and with BU72, two separate featurizations were computed by finding all pairwise ligand atom – receptor residue distances that fall within 10.0 A of the initial, active pose. This amounted to ~700 (ligand atom_i_, receptor residuej) pairs for each ligand.

Features were assessed for “importance” by three main metrics – kinetic slowness (measured by tICA timescale), centrality in a transfer entropy (causality) graph, and predictive ability in cross-validated classifiers. The transfer entropy^18^ was calculated using the MDEntropy package^19^ between all pairs of protein-protein contacts and ligand-protein contacts. To reduce the computational complexity, we included only those protein-protein contacts that were components of the first ten Sparse tICA reaction coordinates, and therefore known to be slowly decorrelating and “important” by this kinetic metric. In total, Transfer Entropy was calculated for approximately 500,000 pairs of features for BU72 and for Sufentanil simulations. A graph was then computed in NetworkX^41^ for each condition, with each node a feature and each directed edge weighted by the Transfer Entropy between them. The PageRank^20^ algorithm was then applied using the NetworkX package as well. In addition, we estimated the contribution of a given feaure (residue-residue or ligand-residue contact) on deactivation by computing with Djikstra’s algorithm in NetworkX the shortest 1,000 paths connecting the ligand to the TM6-in feature, and incrementing for each feature the flux (product of edge weights) through that path: This graph is available in the open GraphML format in the supplementary Information section.

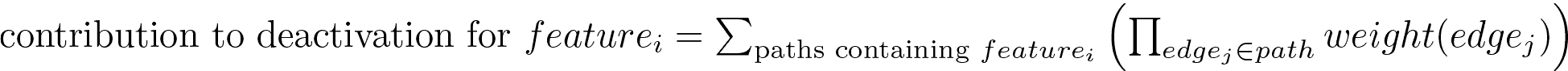

The program Forcefield Toolkit (FFTK) was applied for parameterizing the ligands BU72 and Sufentanil. For parameterization in general and for optimization, charge calculation, and torsion scan in particular, the MP2: 6-31 G^∗^ level of theory and basis set were used in the Gaussian software package, which is a Quantum Mechanics code. To parameterize in a reasonable amount of time, the ligands were fragmented into three sections; torsion scans were performed individually on each fragment as well as on the boundaries between the fragments to yield accurate, whole ligand force fields.

Due to the lack of crystal structures of fentanyl derivatives co-crystallized with μOR, an active-like pose of Sufentanil was derived computationally. First, after removing BU72, Sufentanil was docked into the orthosteric pocket of the active crystal structure with Glide (Schrodinger Inc., New York, NY) with a 25 Angstrom bounding box in the SP setting after first using LigPrep and grid generation tools also included with Schrodinger software packages. Subsequently, the distinct docked poses, which demonstrated significant differences in orientation, were then simulated in NAMD^42^ on the BlueWaters supercomputer. The pose with the lowest average RMSD in simulation was then selected for use in further MD simulation. It is our intention that this approach be replicated for simulating other key ligands of biological systems for which there is no known crystallographic pose.

To more efficiently sample the conformational energy landscape of μOR, two rounds of simulation were conducted. In the initial round, simulations were initiated from the active crystal structure for the unliganded and BU72-bound cases and from the docking- and MD-derivated active-like pose for the Sufentanil-bound case. Then, after approximately 120 μs of simulation for each condition, these “Generation 1” simulations were reseeded with the following procedure. After fitting separate sparse tICA and K-Means models for each condition, those K-Means clusters with the lowest populations were used as the basis for a second round of simulation with randomly reinitiated velocities, termed the “Generation 2” simulations. This strategy is summarized in Extended Data Table 11. MD conformations were deemed inactive/deactivated if they featured a Calpha distance between M161 and V282 of less than 9.0 Angstroms and a closest heavy-atom distance between C159 and Y326 of greater than 10.0 Angstroms.

Visualization was conducted with the MacPyMOL package (Schrödinger, Inc., New York, New York), VMD^43^, and UCSF Chimera^44^.

## Acknowledgements

The authors express their appreciation to Brooke E. Husic, Mohammad M. Sultan, Susruta Majumdar, Bharath Ramsundar, Nathaniel Stanley, Ariana Peck, Danielle J. Marshak, and Samuel Hertig for their insightful input while crafting this manuscript. Outside of Stanford, we thank Robert Abel and colleagues at Schrödinger, Inc. and John Chodera and lab at Sloan-Kettering Institute for insightful input on this work. We thank Samuel Hertig for his assistance in the visual presentation of figures and movies. We also would like to thank Killian Cavalotti, Jimmy Wu, and Stephane Thiell for their indispensable computing support. We thank NIH training grant T32 GM08294 and NIH grant 1171245-309-PADPO for funding, as well as acknowledge the use of Blue Waters at the National Center for Supercomputing Applications and the University of Illinois.

## Author Contributions

E.N.F. and A.B.F. contributed equally to this work. E.N.F. and V.S.P. conceived, and V.S.P. supervised, this project. A.B.F. and E.N.F. set up simulation systems. A.B.F. parameterized BU72 and Sufentanil in Gaussian for the CHARMM force field. E.N.F. generated the prospective docking poses of, and A.B.F. conducted (with NAMD) and analyzed the preliminary simulations of Sufentanil in various poses. E.N.F. conducted all other simulation (listed in E.D. Table 11) in AMBER on XStream. E.N.F. wrote the Python-based code used in analysis of these simulations. C.X.H. and E.N.F. conducted transfer entropy analysis.

## Footnotes

i Notably, the PageRank algorithm^45^ finds that the I256^5.62^–L275^6.30^ distance, a proxy for TM6-TM5 packing (cf. E.D. Table 2, 5), are among the most central nodes in the inter-residue causality (transfer entropy) network of μOR.

